# Directed Functional and Structural Connectivity in a Large-Scale Model for the Mouse Cortex

**DOI:** 10.1101/2021.01.28.428656

**Authors:** Ronaldo V. Nunes, Marcelo Bussotti Reyes, Jorge F. Mejias, Raphael Y. de Camargo

**Affiliations:** Center for Mathematics, Computing and Cognition, Universidade Federal do ABC, São Bernardo do Campo, Brazil; Swammerdam Institute for Life Sciences, University of Amsterdam, Amsterdam, The Netherlands

## Abstract

Inferring the structural connectivity from electrophysiological measurements is a fundamental challenge in systems neuroscience. Directed functional connectivity measures, such as the Generalized Partial Directed Coherence (GPDC), provide estimates of the causal influence between areas. However, the relation between causality estimates and structural connectivity is still not clear. We analyzed this problem by evaluating the effectiveness of GPDC to estimate the connectivity of a ground-truth, data-constrained computational model of a large-scale network model of the mouse cortex. The model contains 19 cortical areas comprised of spiking neurons, with areas connected by long-range projections with weights obtained from a tract-tracing cortical connectome. We show that GPDC values provide a reasonable estimate of structural connectivity, with an average Pearson correlation over simulations of 0.74. Moreover, even in a typical electrophysiological recording scenario containing five areas, the mean correlation was above 0.6. These results suggest that it may be possible to empirically estimate structural connectivity from functional connectivity even when detailed whole-brain recordings are not achievable.

## 1 Introduction

The communication between brain regions is often analyzed using structural and functional connectivity [1]. The former refers to anatomical connections between brain regions generally quantified using tracer injections or diffusion magnetic resonance imaging [2]. The map of these connections is called “connectome” [3]. Network measures are usually used to analyze the connectome, whereas nodes represent brain regions and edges refer to axonal projections [4, 5]. Functional connectivity estimates brain communication from statistical relations between recorded brain signals [6, 1]. Particularly, directed functional connectivity methods use the concept of causality to infer both the intensity and the direction of the connections between brain regions [7]. Even though there is some association between structural and functional connectivity, the relationship between them is not straightforward [1]. While the former is practically static and compose the map of possible pathways for information flow between brain regions, the latter changes continuously and depends, for example, on the dynamical states of brain regions, noise, and strength of structural connections [8].

During electrophysiological procedures, researchers typically record brain signals using electrodes positioned in different depths of brain regions. Even with the improvement in technologies for recording signals, it is usually possible to record signals only from a few areas compared to the number of sources of activity in the brain [9, 10, 11]. Thus, the functional connectivity analysis presents a problem because many unrecorded regions may indirectly influence other regions as common inputs [7, 12, 6]. Therefore, the comparison between structural and functional connectivity becomes more complicated since spurious inferred causality relations can lead to misinterpretations of electrophysiological data.

Previous simulation studies evaluated the relation between directed functional connectivity and structural connections [8, 13, 14, 15]. However, most of these studies used either autoregressive [16] or rate-based models [13] for the dynamics of each cortical area. These studies provided essential steps towards evaluating the reliability of causality measures. However, the time series obtained from autoregressive and rate models are distant from electrophysiological signals obtained in experimental laboratory conditions. Using spiking models, we can capture the dynamic of neuronal networks while generating simulated LFP signals from the synaptic currents. Also, most studies do not consider the impact of accessing only part of the activity in the brain.

In this work, we investigate the relationship between directed functional connectivity and structural connectivity in a large-scale network model of the cortex, derived from a cortical connectome of the mouse obtained using tracer injections [17]. We used GPDC, a frequency-domain method based on multivariate vector autoregressive models, which provides estimates of directed functional connectivity [18, 19]. The mean correlation between FLN and GPDC remained high (*r* > 0.6) even when only a few cortical areas were considered in the GPDC calculation, indicating that this causality measure provides reliable results in typical experimental conditions in which only recordings from a subset of areas are available.

## 2 Methods

### 2.1 Neuron Model

We modeled the neurons using a single-compartment Hodgkin–Huxley-type model, where the membrane potential of the *i*-th neuron described by,

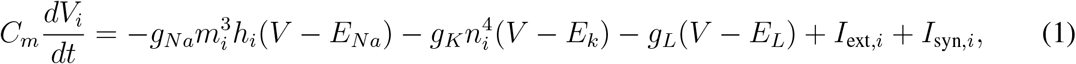

the membrane capacitance *C_m_* is 0.50 nF (0.25 nF) for excitatory (inhibitory) neurons. The maximal conductances values were *g_Na_* = 12.5 *μ*S, *g_K_* = 4.74 *μ*S and *g_L_* = 0.025 *μ*S. The reversal potentials *E_Na_* = 40 mV, *E_K_* = −80 mV, and *E_L_* = −65 mV correspond to the sodium, potassium and leakage channel, respectively [20]. The dynamics of the voltage-gated ion channels are described by activation and inactivation variables *m*, *n*, and *h*, where *m* and *n* accounts for the dynamics of Na channels and *h* for K channels. The probability that an ion channel is open evolves according to a set of ordinary differential equations [21],

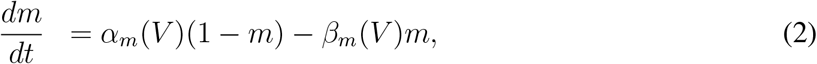

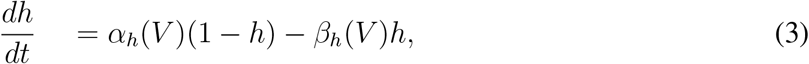

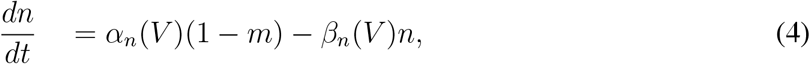

where,

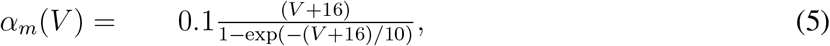

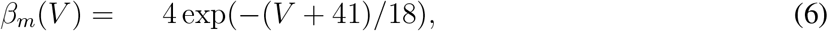

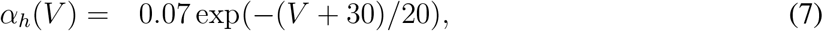

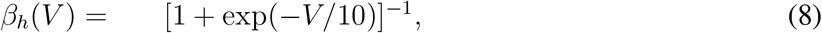

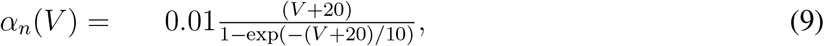

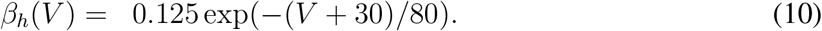

The parameters used in this neuron model were previously reported and applied in some studies that modeled cortical neuronal populations [22, 21, 23].

### 2.2 Spiking neuronal population model

Each spiking neuronal population was composed of 2000 neurons, 1600 excitatory and 400 inhibitory. Connections between neurons within each spiking neuronal population are random with connection probability *p*_intra_ = 10%. The synaptic current *I*_syn_ that arrives to postsynaptic neuron *i* is modeled by,

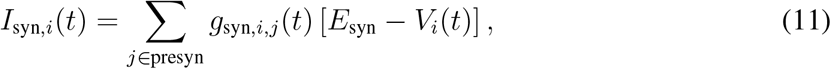

where the index *j* represent a presynaptic neuron connected to neuron *i* and the sum over *j* accounts for all the synapses that impinge on neuron *i*. *E*_syn_ is the synaptic reversal potential which is 0 mV for excitatory and −70 mV for inhibitory synapses. The dynamics of synaptic conductance *g*_syn,*i,j*_ is described by an exponential function as follows [24],

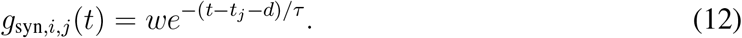

The characteristic decay time *τ* is 2 ms and 8 ms for excitatory and inhibitory synapses, respectively. When a presynaptic neuron *j* fires a spike at time *t_j_*, *g*_syn,*i,j*_ is incremented by a synaptic weight *w* after the axonal delay *d*, which was set as 1 ms for all intra-areal connections [21]. The value of w depends on the excitatory/inhibitory nature of the presynaptic and postsynaptic neurons. Furthermore, all neurons receive a background input given by a heterogeneous Poisson-process spiking activity with a rate of 7.3 kHz [21]. The background input acts as an excitatory synaptic current. To add heterogeneity in our model, all synaptic weights w for recurrent connections and background input were taken from a Gaussian distribution (Table 1).

**Table 1:** Synaptic weights for intra-areal connections. Mean synaptic weight 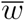 and standard deviation *σ_ω_* for all possible synapses. E, I, and Input represent excitatory neurons, inhibitory neurons, and external input, respectively. The arrow indicates the direction of the connection.

| Synapses | $\bar{w}$ (nS) | $\sigma_w$ (nS) |
| --- | --- | --- |
| $E \rightarrow E$ | 2.5 | 1.0 |
| $E \rightarrow I$ | 2.5 | 1.0 |
| $I \rightarrow E$ | 240 | 10 |
| $I \rightarrow I$ | 240 | 10 |
| $Input \rightarrow E$ | 3.2 | 1.0 |
| $Input \rightarrow I$ | 3.2 | 1.0 |

### 2.3 Mouse large-scale cortical network

The mouse cortex’s large-scale network model is composed of 19 areas where a spiking neuronal population models each area with long-range and recurrent synapses. Parameters related to recurrent synapses were described in the previous session. Neurons from different areas are randomly connected with probability *p*_inter_ = 5%. The synaptic weights between cortical areas are based on the recently published anatomical connectivity dataset for the mouse cortex [17] obtained by retrograde tracer injections [25].

This technique consists in injecting a tracer that flows from the target synapses to the cell bodies, allowing to identify neurons projecting to the target area. The Fraction of Labeled Neurons (FLN) was measured as the ratio of the number of labeled neurons in a source area to the total quantity of labeled neurons in all source areas, where labeled neurons considered are extrinsic to the injected area [25, 26, 27]. We defined the edge measure FLN_*ij*_, as the number of neurons projecting from area *j* to area *i*, divided by the number of neurons projecting to area *i* from all the areas except *i* [28, 29]. The synaptic weights for directed long-range connections are the FLNs scaled by the global scaling parameters *μ*_E_ = 50 and *μ*_I_ = 25,

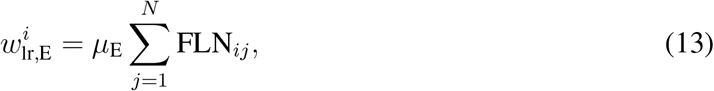

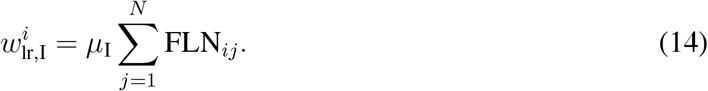

Long-range connections are excitatory, targeting either excitatory or inhibitory neurons with synaptic weight, 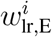, and 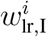, respectively. The index *j* represents the source area, *i* the target area, and *N*represents the resistance of a typical electrode is the total number of simulated cortical areas. The axonal delay for long-range connections is given by the ratio between the inter-areal anatomical distance estimates between cortical areas and the conduction speed set as 3.5 m/s [30].

### 2.4 LFP signal

We computed the local field potential (LFP) signal as a sum of the currents’ absolute values acting upon excitatory neurons in a spiking neuronal population [31,32]. Thus, for a cortical area in our model, the LFP signal will be given by,

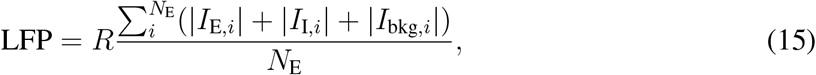

*I*_E,*i*_ accounts for both the local (within population) and global (inter-areal projections) excitatory synaptic currents, while *I*_I,*i*_ corresponds to the local inhibitory current. *I*_bkg,*i*_ is the synaptic current related to the background Poisson input. *R* represents the resistance of a typical electrode used for extracellular measurements, here chosen to be 1 MΩ [21]. NE is the number of excitatory neurons in each neuronal population.

The mean was subtracted from the simulated LFP signal. The resultant signal was filtered using a 1 kHz low-pass filter to avoid aliasing and downsampled to 1 kHz.

### 2.5 Generalized partial directed coherence

Generalized partial directed coherence (GPDC) is a frequency-domain method of directed functional connectivity established on multivariate vector autoregressive (MVAR) model [18]. The MVAR model for a set x(*t*) = [*x*_1_(*t*) … *x_N_*(*t*)]^*T*^ of simultaneously observed time series is defined as:

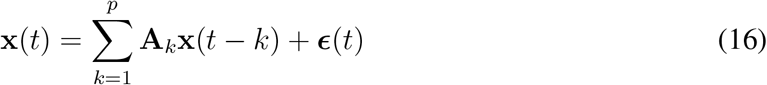

where *p* is the MVAR model order. Akare coefficient matrices in which the element *A_ij,k_* define the effect of *X_j_* (*t* − *k*) on *x_i_*(*t*), where *k* is the time lag. The term *ε*(*t*) is a vector of *N* white noises with covariance matrix Σ. The GPDC from the time series *x_j_* to the time series *x_i_* at frequency λ is defined as,

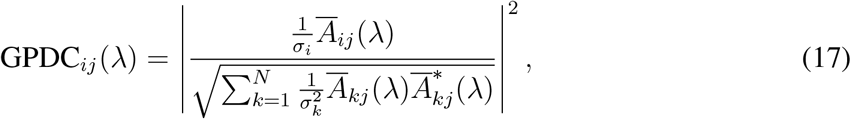

where

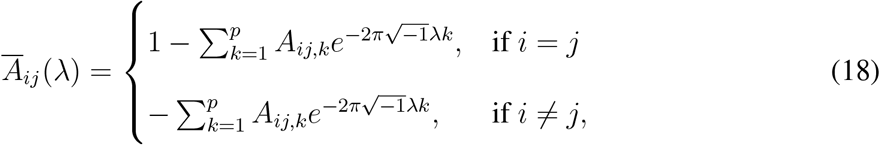

and 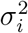 refers to the variance of white noise *ε*_*i*_(*t*) [18]. λ is a normalized frequency where |λ| ≤ 0.5 so that λ = 0.5 means one half of the sampling rate *f_s_* [19]. The MVAR model was estimated by the method of ordinary least squares (OLS) [33]. We used Akaike’s Information Criterion (AIC) to select model order (Supplementary Equation S1), choosing the order *p* ≤ 50 that had the minimum AIC (Supplementary Figure S6) value.

GPDC has values in the range from 0 to 1 and is invariant to scale, so the normalization of time series is unnecessary [19, 18]. Similar to other (directed) functional connectivity measures, unrecorded time series can lead to spurious estimates. Therefore, the reliability of estimates depends on the number of time series included in the estimates. For all analysis we used the peak GPDC value over all frequencies 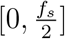.

### 2.6 Estimated activity

The activity flow mapping measures the propagation of neural activity by estimating the activation of a target region. It is defined as the sum of the activity in each source region multiplied by the functional connectivity with the target region [34]. We adapted the idea of activity flow by defining two measures of estimated activity arriving in a cortical area *i* mediated by pathways of structural connectivity (FLNs) and directed functional connectivity (GPDC peak),

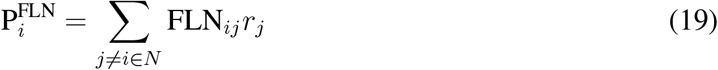

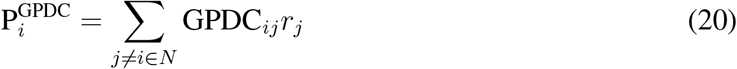

where FLN_*ij*_ is the FLN from area *j* to area *i*, *r_j_* is the firing rate for area *j*, GPDC_*ij*_ is the peak of GPDC from area *j* to area *i*, and N is the total number of simulated cortical areas.

### 2.7 Centrality measure

We computed the nodal in-strength for the mouse cortical connectome. The nodal in-strength for a node *i* is given by

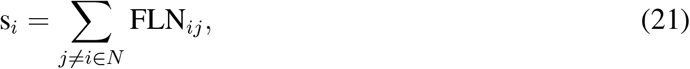

where *j* is the source area, *i* the target area, and *N* the total number of simulated cortical areas [35].

### 2.8 Numerical simulations

All simulations were performed using the simulator Brian2 [36] applying the exponential Euler method [37] to integrate the differential equations with an integration step of 0.1 ms. Each simulation was 30 s long, generating sufficient data points to apply GPDC on the simulated LFP signals [38].

## 3 Results

The large-scale network model of the mouse cortex contains 19 spiking neural populations with recurrent connections and excitatory long-range connections between populations, constrained by the directed and weighted structural connectome (Figure 1A and Figure 1B). The dynamical behavior of each simulated cortical area is predominantly asynchronous with transient spike synchronization [39, 40] (Figure 1C), with the typical power spectral density (PSD) of LFP signals displaying a peak in the gamma band (Figure 1D and 1E) [41]. The firing rate of inhibitory neurons is 4.74 ± 0.11, higher than the excitatory neurons rate of 3.64 ± 0.42 (Figure 1F). Differences in population behavior are mostly due to inputs from other areas since we sample their parameters from the same distributions.

**Figure 1:**
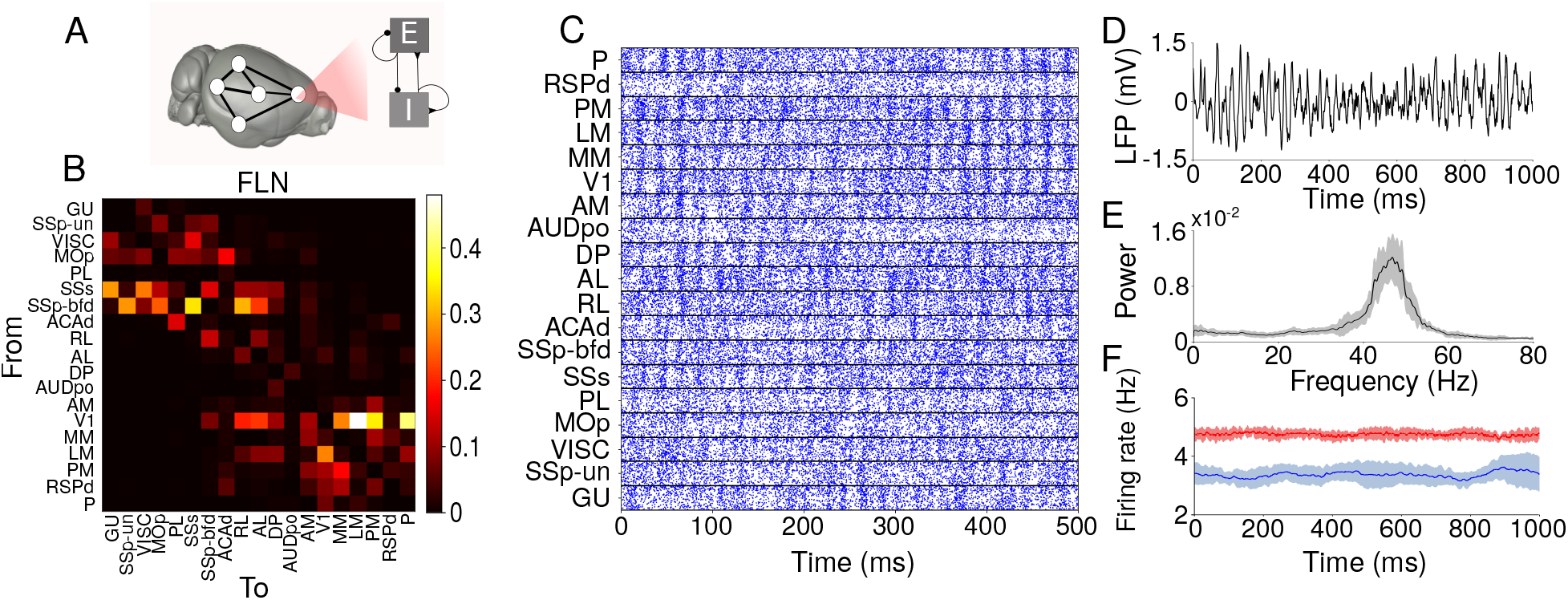
Large-scale cortical network. (A) Local neuronal population where E and I are populations of spiking neurons [42, 43]. (B) Map of structural connectivity given by the FLNs. These values define the strength of long-range projections in the large-scale network model. (C) Raster plot of 500*ms* of activity for each cortical area. (D) Simulated LFP signal for an area in the large-scale network model. (E) Power spectral density for simulated LFP signal for one area. The continuous black line corresponds to the average over ten simulations, and the gray shaded area delimits its standard deviation. (F) Firing-rate for excitatory (blue) and inhibitory (red) populations computed using a sliding window of 100*ms*. The continuous line corresponds to an average firing rate over ten simulations, and the shaded area is the standard deviation. To exemplify, we used data from area MOp in (D), (E), and (F).

We first compared the FLN values to the average GPDC over ten simulations of the model. Most medium to strong connections from the structural connectome were also captured by the directed functional connectivity (Figure 2A and Figure 2B). We used the GPDC largest value (peak), but other approaches such as the average of GPDC over frequencies and area under the GPDC curve (Supplementary Figure S1) produced similar results.

**Figure 2:**
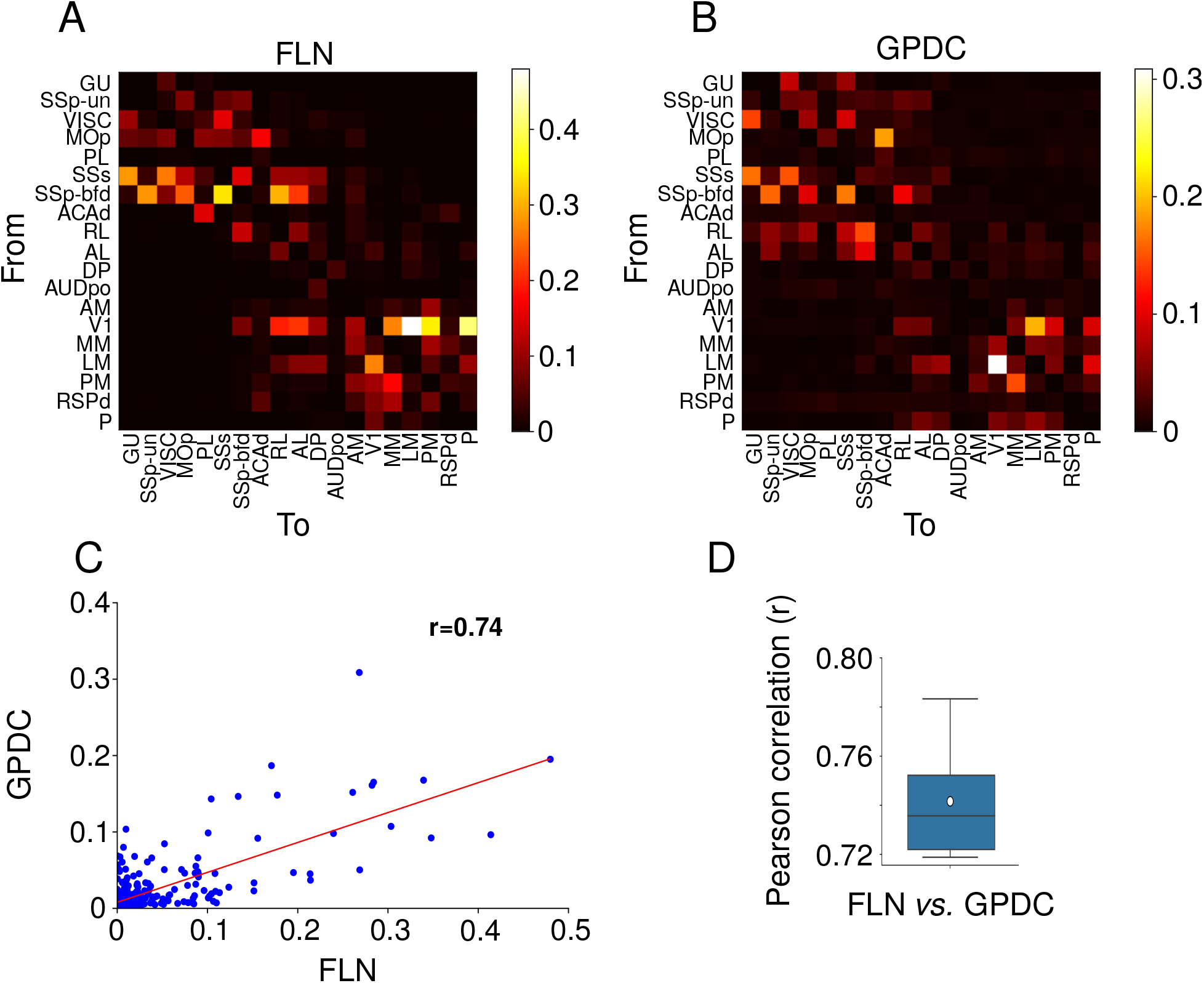
Relation between structural and directed functional connectivity. (A) Map of structural connectivity given by the FLNs. (B) Map of directed functional connectivity given by GPDC peaks for one simulation. GPDC from a cortical area to itself was set as 0. (C) Scatter plot of FLNs *versus* GPDC peak for one simulation. The red line corresponds to the linear fit. The Pearson correlation between FLNs and GPDC is 0.74. (D) Box plot showing the distribution of Pearson correlation between FLN and GPDC for ten simulations. The white circle represents the average Pearson correlation over ten simulations which is 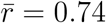.

Although the graph density of the structural connectome is 97% [17], most structural connections are weak, which leads to a prevalence of weak average GPDC values. Weak structural connections is a characteristic shared by connectomes from different mammals, with FLNs varying by several orders of magnitude, log-normally distributed [25, 26, 17, 44]. To evaluate the relation between structural and directed functional connectivity, we plotted GPDC values from ten simulations against FLNs and fitted a linear model, obtaining the Pearson correlation *r* (Figure 2C). The scatter plot presents most points close to the origin due to the predominance of small values for the GPDC and FLN. The average Pearson correlation between FLN and GPDC is 0.74. We also verified that the average correlation between GPDC and FLN over bootstrap samples of 80 randomly chosen edges is 0.74 (Supplementary Figure S2). This correlation level is close to those obtained by other works that analyzed different structural connectomes using functional connectivity applied to empirical data (*r* ≈ 0.79 [45]) or firing rate models (*r* ≈ 0.73 [46]).

The centrality of the cortical area seems to influence the variability of GPDC estimates over simulations. The variability of directed functional connectivity was measured by the coefficient of variation of GPDC (Figure 3A). The centrality, measured by the nodal in-strength (i.e., the sum of inward FLNs to a cortical area) (Figure 3B), is positively correlated (*r* = 0.64) to the sum of the coefficients of variation (CVs) of the connections emerging from that area (source) (Figure 3C). When the cortical area is considered the target of directed functional connectivity, the correlation with nodal in-strength is negative (*r* = −0.52). We performed the same analysis correlating the sum of coefficient of variation with eigenvector centrality (Supplementary Figure S3), and we obtained the same relationship, but with smaller Pearson correlation coefficients (*r* = 0.59 and *r* = −0.44). We should note that in both cases (source and target), the actual variability (standard deviation) increases with larger nodal in-strength values (Supplementary Figure S4).

**Figure 3:**
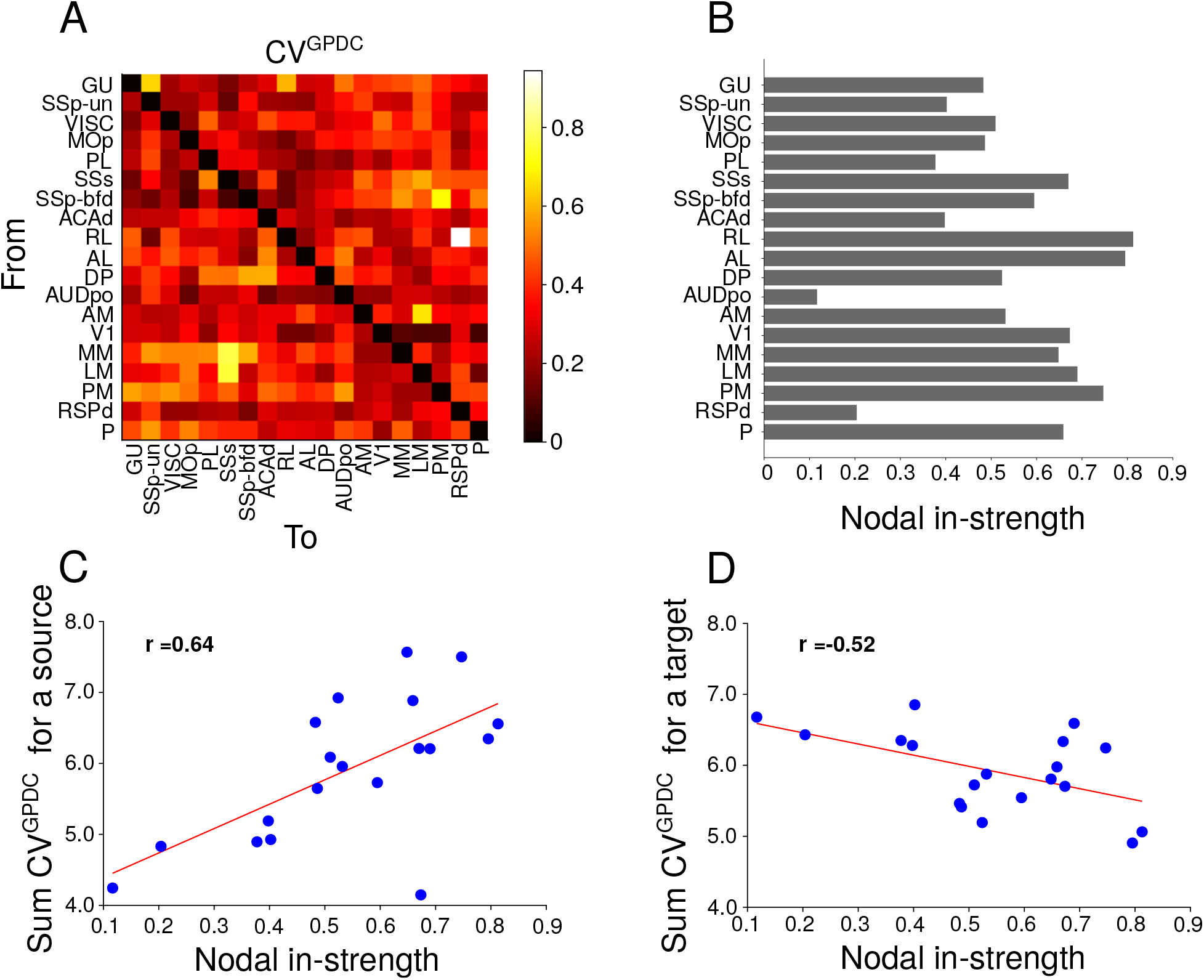
Relationship between nodal in-strength and varibility of GPDC. A) Coefficient of variation for GPDC (CV^GPDC^). B) Nodal in-strength for all cortical areas. C) Sum of CVGPDC for a source (Sum of columns in A) vs. nodal in-strength. D) Sum of CV^GPDC^ for a target (Sum of rows in A) *vs*. nodal in-strength.

We also investigated the relationship between the firing rate in a cortical area and the estimated activity that is arriving at this cortical area mediated by structural or directed functional connectivity pathways. The propagation of activity in the cortex is constrained by direct anatomical connections between areas and indirect paths [47], with the propagation of activity occurring mainly through the strongest long-range projections [28]. The estimated activity mediated by FLNs is strongly correlated to the target areas’ firing rate (Figure 4A), while the correlation of estimated activity mediated by GPDCs and firing rates was 0.54 (Figure 4B). This indicates that GPDC estimates can be used to infer the propagation pathways, although less reliably than when using FLN values directly.

**Figure 4:**
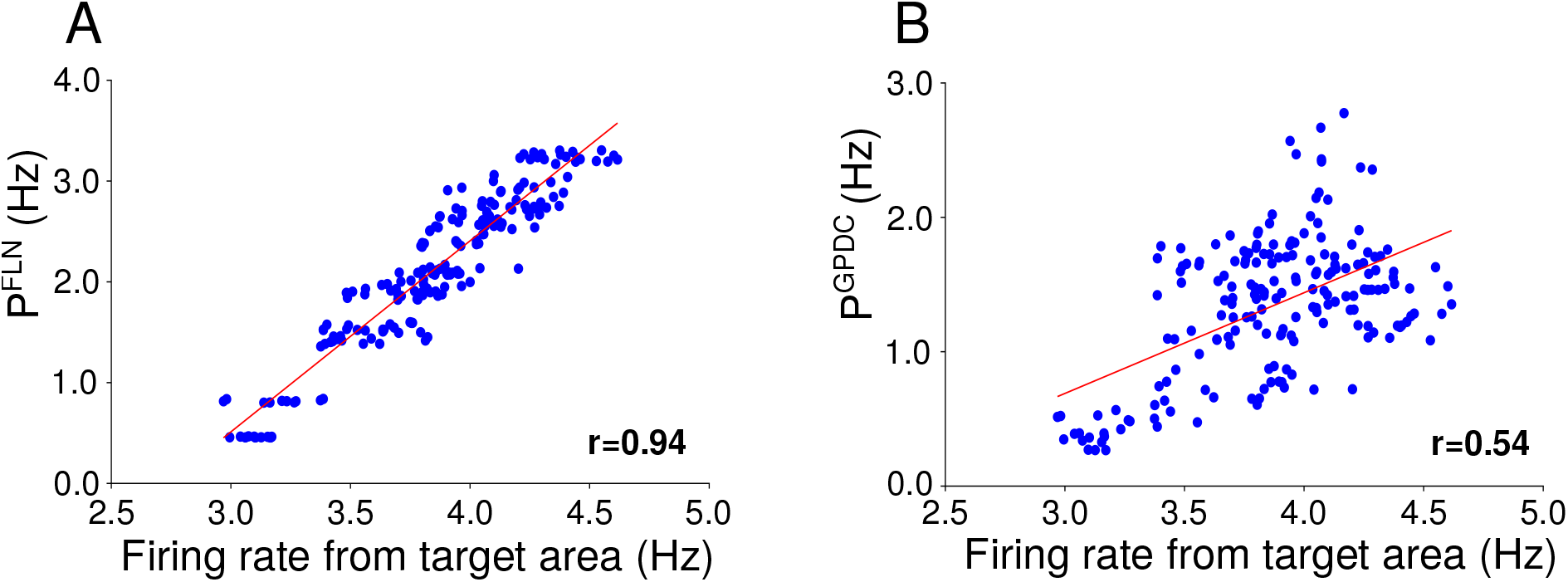
Estimated activity through structural and directed functional pathways. (A) Estimated activity mediated by structural connectivity versus firing-rate for target areas (*r* = 0.94). (B) Estimated activity mediated by directed functional connectivity versus firing-rate for target areas (*r* = 0.54). Red lines are linear fits.

We analyzed the behavior of GPDC estimates when considering a reduced number of areas, reproducing typical experimental setups. We considered a visual and a frontoparietal cluster, each containing 7 cortical areas [17] (Figure 5A). We evaluated the distribution of correlation between FLN and GPDC when GPDC estimates between all areas of each cluster are conditioned on the whole connectome, conditioned on the areas in each cluster, and using only pairwise (bivariate) estimates (Figures 5B and 5C). This analysis simulates the situations where an electrophysiologist only has information from a single cluster of cortical areas or a pair of areas. The highest correlations between the GPDC and FLN occurred when we conditioned GPDC to the whole connectome, followed by GPDC conditioned to the cluster area, and pairwise GPDC. Also, the correlation for the frontoparietal cluster was higher than for the visual cluster in all scenarios.

**Figure 5:**
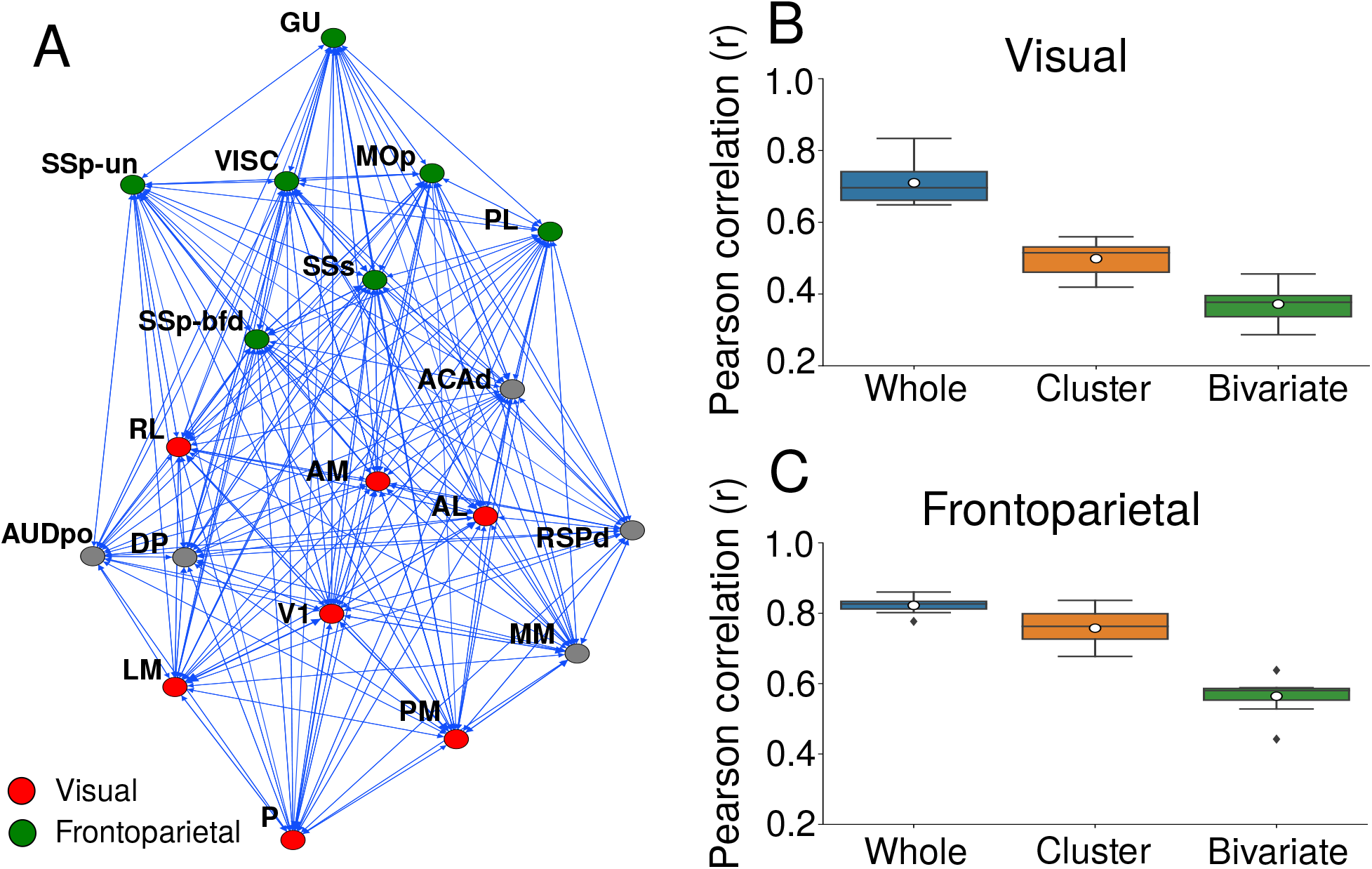
Correlation between FLN and GPDC for the visual and frontoparietal clusters. (A) Graph representing the mouse cortical connectome. Nodes represent cortical areas and edges, directed long-range projections between them. Green nodes are cortical areas belonging to the frontoparietal cluster. Red nodes are cortical areas belonging to the visual cluster. Each cluster contains 7 cortical areas. Box plot of Pearson correlation between FLN and GPDC for the visual cluster (B) and frontoparietal cluster (C). GPDC was computed considering the whole connectome (blue box), only the cluster (orange box), and pairwise (green box).

We extended the analysis to evaluate the effect of cluster size on GPDC correlation to FLN. We used cluster sizes ranging from 2 to 15 areas. We created 150 random clusters sampled from all areas in the connectome for each cluster size and computed the Pearson correlation for the GPDC (A) conditioned on the whole connectome, (B) conditioned on the cluster areas, and (C) evaluated using pairwise data. For cases (A) and (B), the Pearson correlation increases, and the standard variation decreases as we increase the cluster size (Figure 6), showing that it is advantageous to include more areas in the GPDC calculation. Surprisingly, the correlation between structural and directed functional connectivity when using simulated signals from a few cortical areas (blue dots) is similar to using signals from the whole cortex (black dots), with most points showing statistically different results. The bivariate GPDC (Figure 6) had a statistically significant lower average Pearson correlation for all cluster sizes with four or more areas, indicating that these measures are affected by interference from ignored signals.

**Figure 6:**
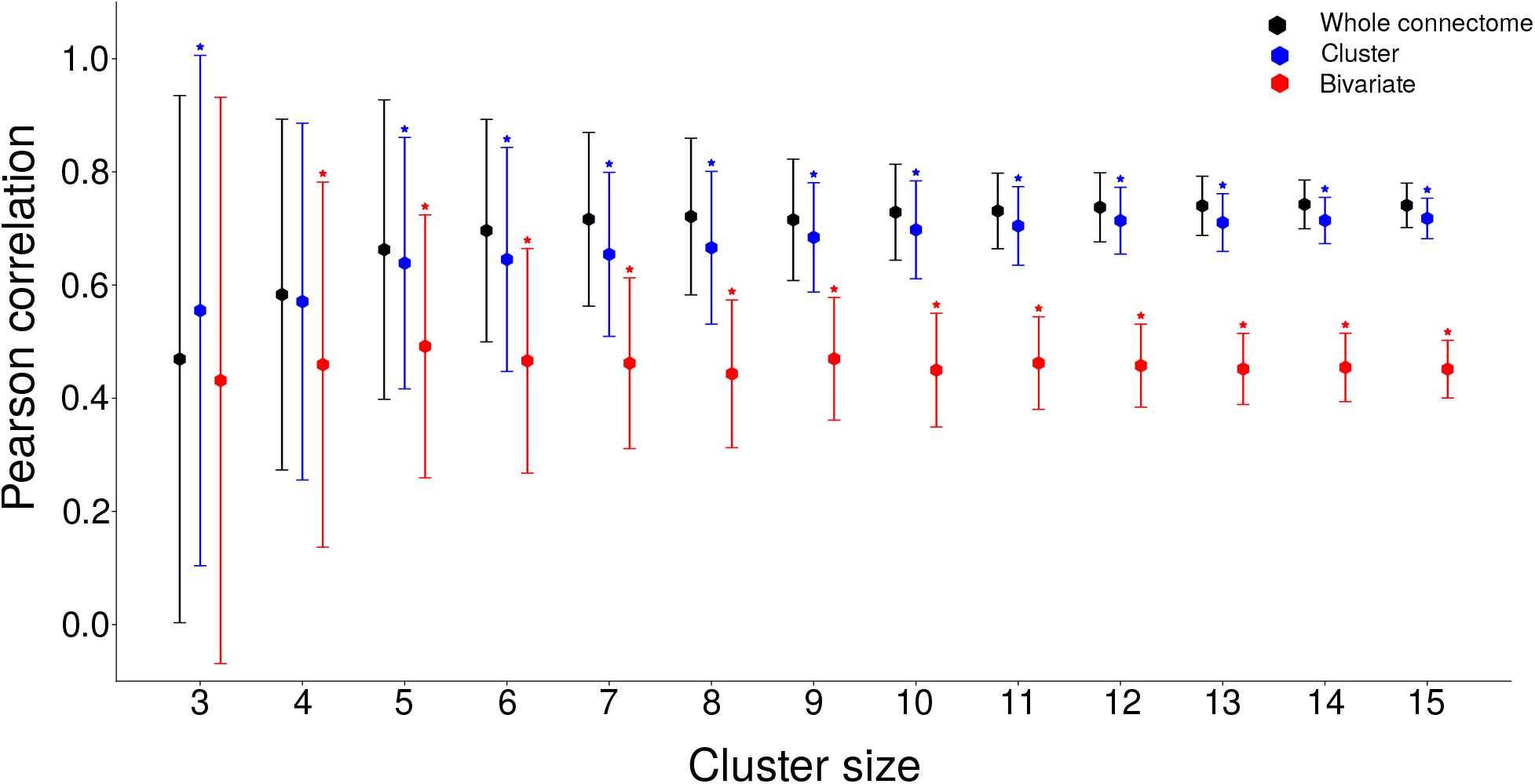
textbfPearson correlations between FLN and GPDC for the different cluster sizes with randomly chosen areas. The graph shows the average (dots) and error bars of the GPDC conditioned to the whole connectome (in black), to areas in the cluster (in blue), and to pairwise signals (in red). The error bar is the standard deviation of Pearson correlation considering values for all randomly chosen areas and all simulations. Stars represent statistically different averages when compared to the whole connectome (Holm-Bonferroni corrected Welch t-test).

## 4 Discussion

Our results shed light on the relationship between structural and directed functional connectivity in circumstances similar to those faced by electrophysiologists. They indicate that the reliability of directed functional connectivity estimates and their relationship with structural connectivity depends on the number of areas considered. Nevertheless, the GPDC conditioned on few cortical areas had similar results to the GPDC conditioned on all areas, providing evidence that it is possible to obtain a reasonable relationship between structural and directed functional connectivity in electrophysiological experiments even with signals from few areas.

Previous studies evaluated the relationship between structural and functional network connectivity strength on electrophysiological data [48], with some using undirected functional connectivity measures [49, 50]. But in electrophysiology studies, researchers do not have access to signals from unrecorded areas and have only estimates of structural strengths from tracers. Using large-scale network models solves this problem, as the researcher has access to all variables in the system, allowing a better understanding of the obtained functional connectivity results.

The relationships between structural and functional connectivity have been largely unexplored through large-scale network models [51], and the existing models use neural mass descriptions (rate models) to describe each area’s activity [52, 53]. However, information propagated between brain regions can be characterized not only by the rate code but also by the temporal code [54, 55, 56, 57,58], and hypotheses are pointing to spike-timing and spike coherence as essential components of cortical communication [59, 39, 60]. Spiking neuronal populations have richer dynamical behaviors than rate models and better resemble cortical activity, through spiking neuronal networks it is possible to investigate the consequences of spike synchronization [39], model different approaches for the propagation of information [28, 56], and generate simulated LFP signals from the synaptic currents, which better resemble biological LFP signal [31]. Moreover, our obtained correlations are in the same range as the studies using more complex electrophysiological data [48].

The centrality of a cortical area affects the variability of GPDC estimates in different ways when such area is examined as the target or source of functional connections. Strong functional connectivity generally occurs between areas with direct structural connections [53], and network measures applied to structural connections can help predict the resting-state functional connectivity [61]. However, as far as we know, no previous work has indicated that the variability of directed functional connections could be partially explained by centrality measures applied to structural connectivity. We also noticed that synchronization is strongly correlated to the centrality of the node (Supplementary Figure S5). So it is likely that stronger long-range connections targeting an area increase the synchronization of spikes in this area, and the increased synchronization changes the variability in directed functional connectivity. Indeed, it was observed in previous work that synchronization has an important role in directed functional connectivity [39].

The firing rate of cortical areas is explained by the estimated activity flow, as proposed by Cole *et al*. [34]. When using GPDC as an estimate of structural connections, the correlation between actual and estimated activity in the target area decreases to 0.54. This indicates that directed functional connectivity can be used to estimate the activity flow. Although it is less reliable than when using the actual structural connection strengths, researchers may only have access to directed functional measures.

The relationship between structural and directed functional connectivity is the largest when GPDC is conditioned to all areas in the connectome and decreases as we reduce the number of areas. GĂmĂnut et al. identified 6 clusters in the mouse connectome (prefrontal, frontal, parietal, cingulate, temporal, and visual) [17] based on the same approach used to investigate the macaque cortex [62]. We evaluated the relationship between GPDC and FLNs in the visual cluster and in a combination of the prefrontal, frontal and parietal clusters, which we called frontoparietal. We did not use the other clusters, which had a small number of regions. The average correlation was in the range of correlation obtained for random clusters, with *r* = 0.76 for the frontoparietal and *r* = 0.50 for the visual cortex. This indicates that within anatomical clusters the relationship between GPDC and FLNs does not change in relation to randomly selected areas. These results also show that GPDC estimates provide statistical information on structural connections even when considering only a few areas. However, when considering individual connections, there can be large differences between GPDC estimates and actual structural connection strengths.

Our large-scale network model has some limitations. First, modeled neuronal population parameters are drawn from the same distributions with activity in the gamma band range (Figure 1). The activity of cortical areas in mice occurs in multiple frequency ranges [63, 64] and the relationship between structural and functional connectivity depends on the frequency [27]. A second limitation is that we do not model changes in network states, which are known to influence functional connectivity [65]. Some studies in computational neuroscience have already explored multistability and temporal patterns of functional connectivity [66, 67, 68]. Finally, we considered only cortical areas in our large-scale network model, excluding subcortical areas, which have a more complex dynamic [69, 70]. Future studies can overcome these limitations by creating richer spiking network models, with different operating frequencies and evolving neuronal dynamics. These models are difficult to create but would allow one to compare functional connectivity values to structural connection strength in more dynamic settings.

## Supplementary Material

**Figure S1:**
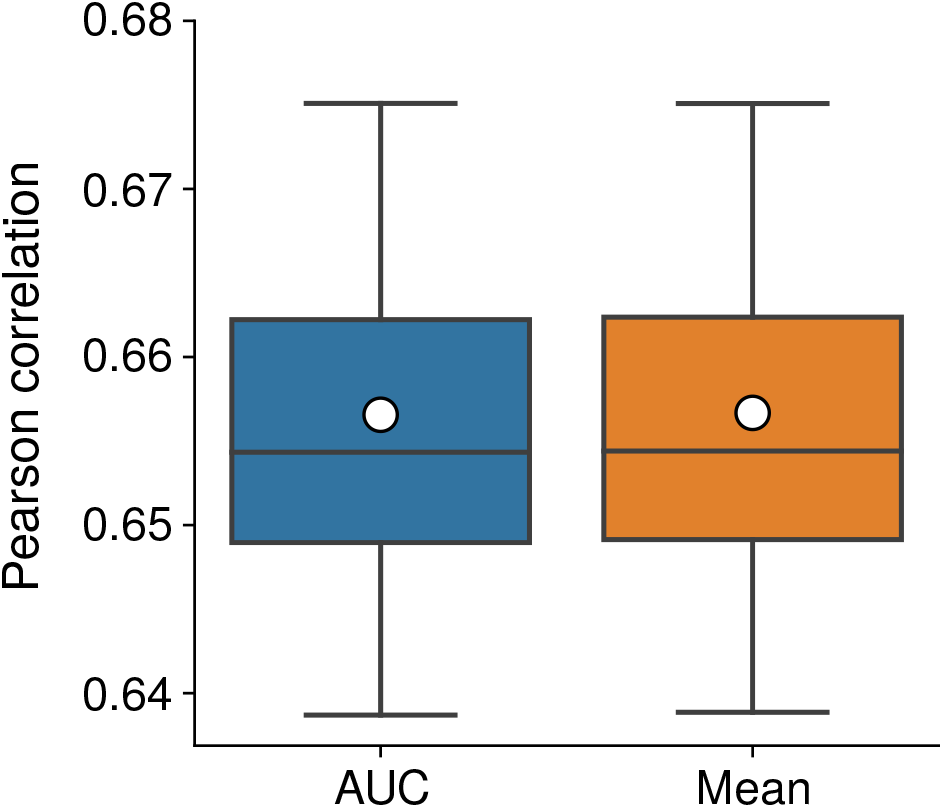
Distribution of correlation between area under curve (AUC) of GPDC estimates or mean of GPDC and FLN. In blue, distribution of Pearson correlations between AUC of GPDC and FLN. The average Pearson correlation is 0.65. In orange, distribution of Pearson correlations between mean of GPDC and FLN. The average Pearson correlation is also 0.65.

**Figure S2:**
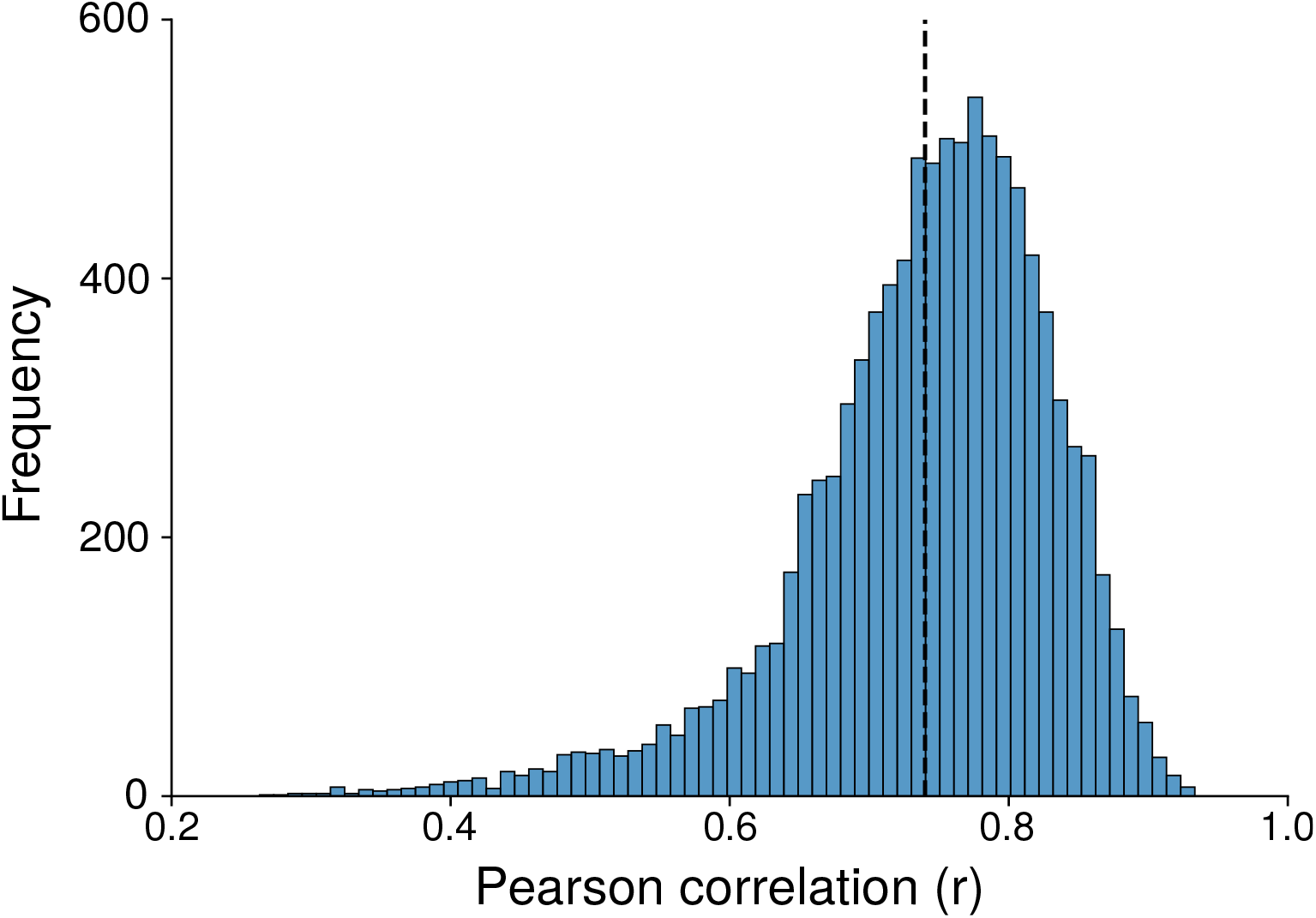
Distribution of correlation between FLN and GPDC for 1000 bootstrap samples of 80 randomly selected edges. In each bootstrap sample, it was computed the correlation for each simulation separately, involving a total of 10000 samples (10 simulation x 1000 bootstrap samples). The dashed line is the mean of the distribution 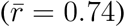.

**Figure S3:**
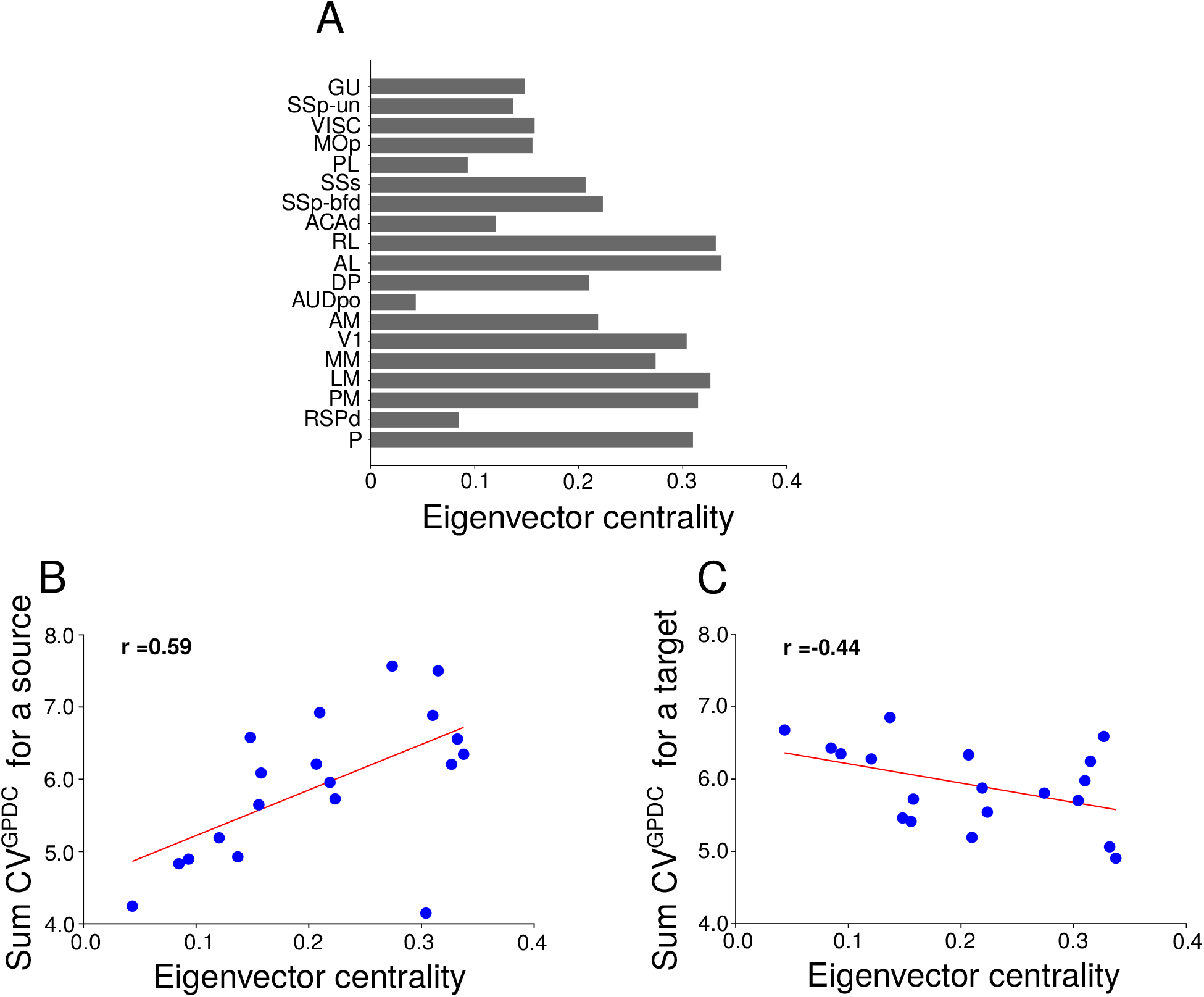
Relationship between eigenvector centrality and varibility of GPDC. A) Eigenvector centrality for all cortical areas. B) Sum of CV^GPDC^ for a source vs. eigenvector centrality. C) Sum of CV^GPDC^ for a target *vs*. eigenvector centrality.

**Figure S4:**
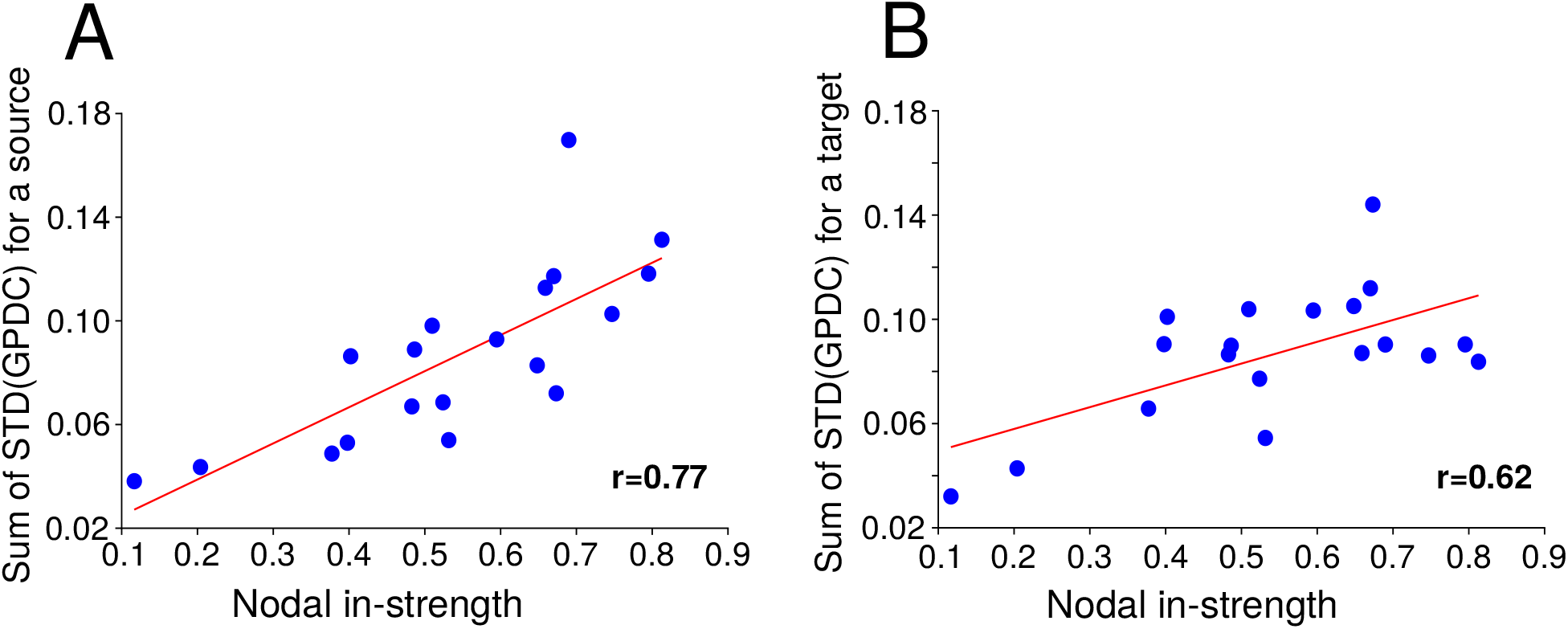
Standard deviation of GPDC and centrality. A) Sum of standard deviation of GPDC for a source *vs*. nodal in-strength. C) Sum of standard deviation of GPDC for a target *vs*. nodal in-strength.

**Figure S5:**
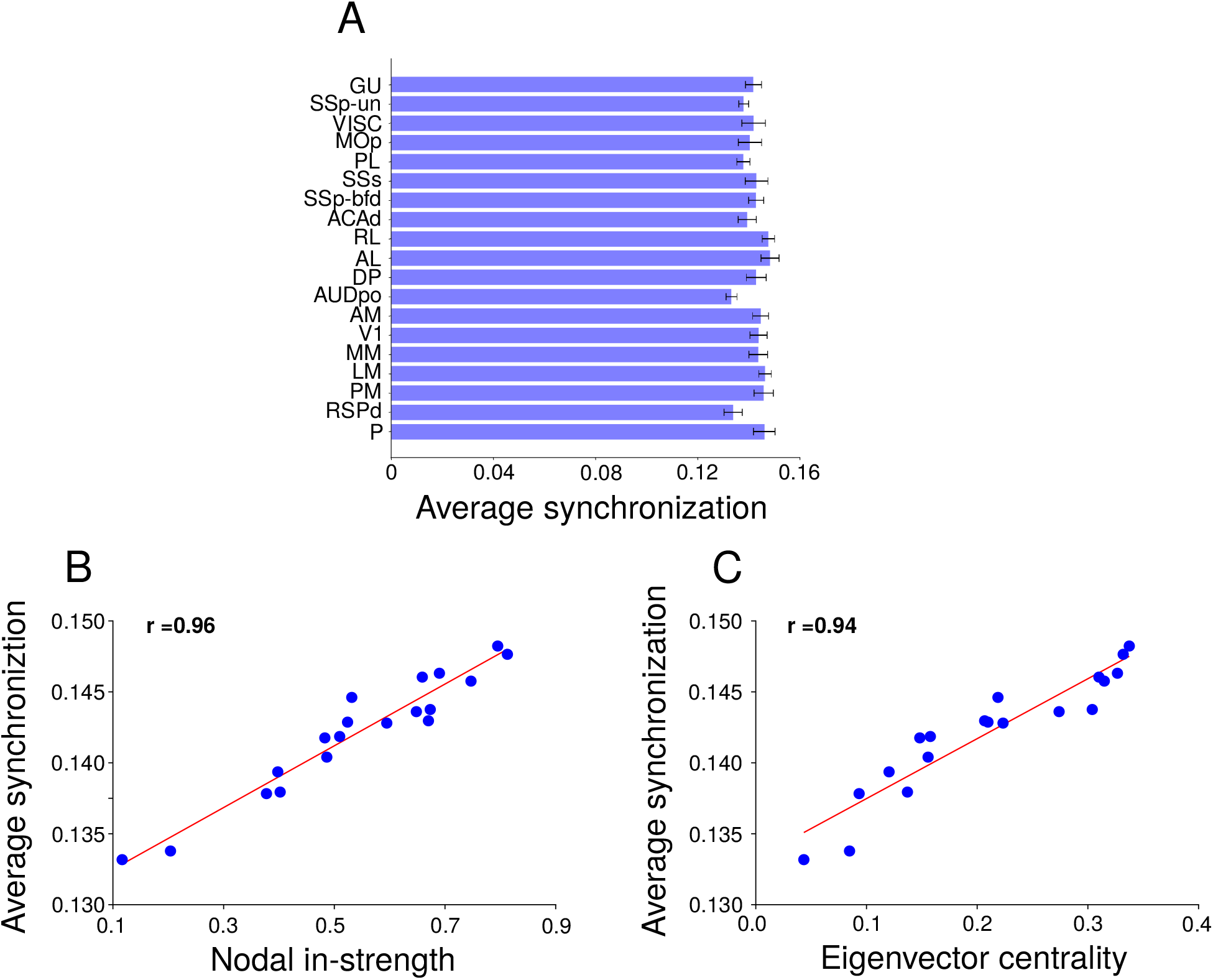
Synchronization *vs*. centrality. A) Average synchronization (average over simulation of average synchronization over time). Bars are standard deviation. B) Average synchronization vs nodal in-strength. C) B) Average synchronization vs eigenvector centrality. Synchronization was obtained using PySpike [71]. Eigenvector centrality was computed using NetworkX [72].

**Table S1:**
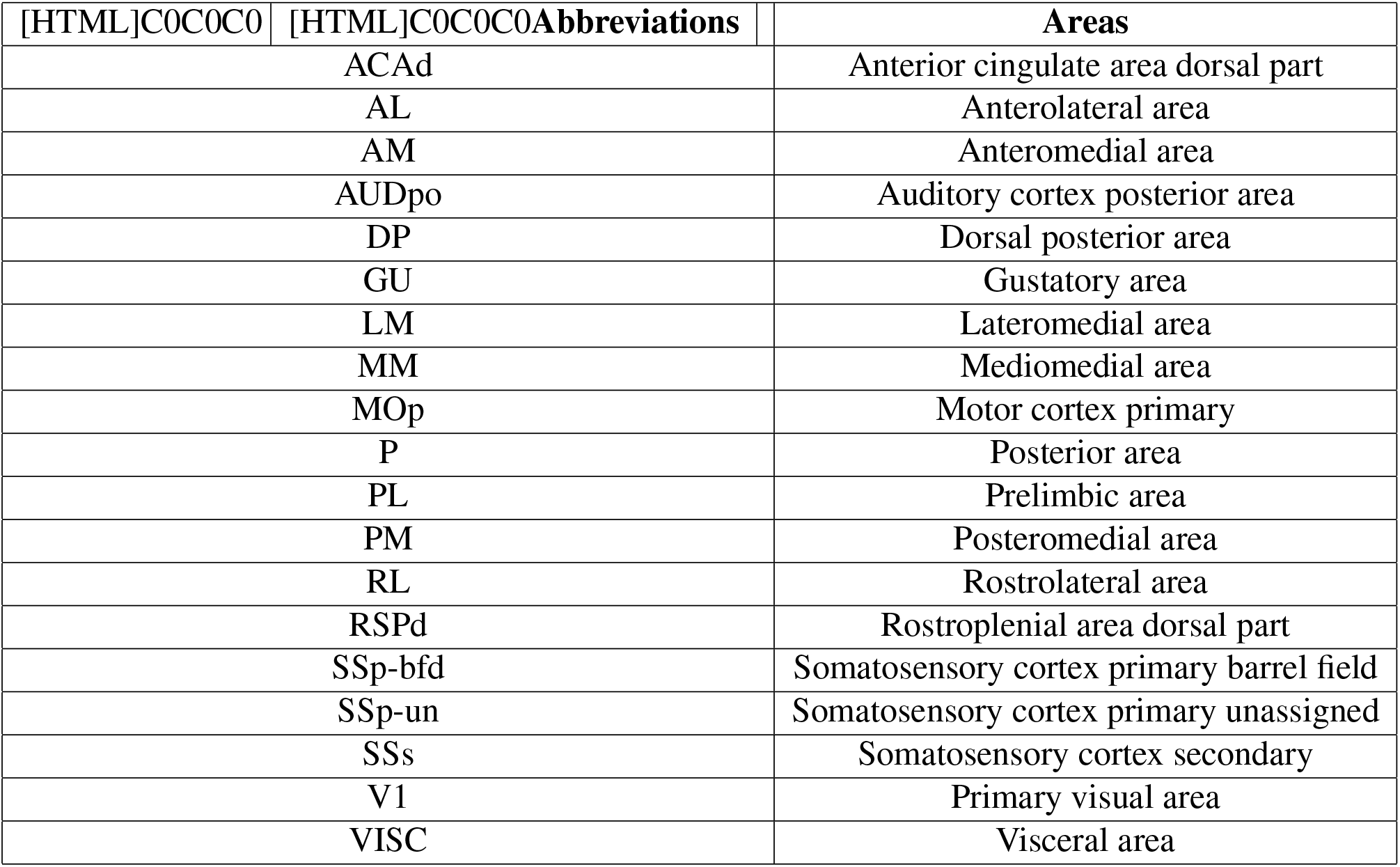
Name of areas in mouse cortical connectome. Adapted from Gămănut et. al., 2018.

### Akaike’s Information Criterion (AIC)

The AIC for order *p* is obtained by

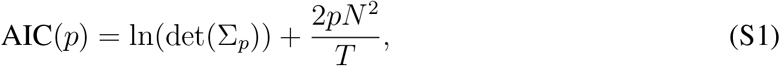

Σ_*p*_ is the covariance matrix of residuals for the model with order *p*, *N* is the number of time-series and *T* is the length of time-series [19].

**Figure S6:**
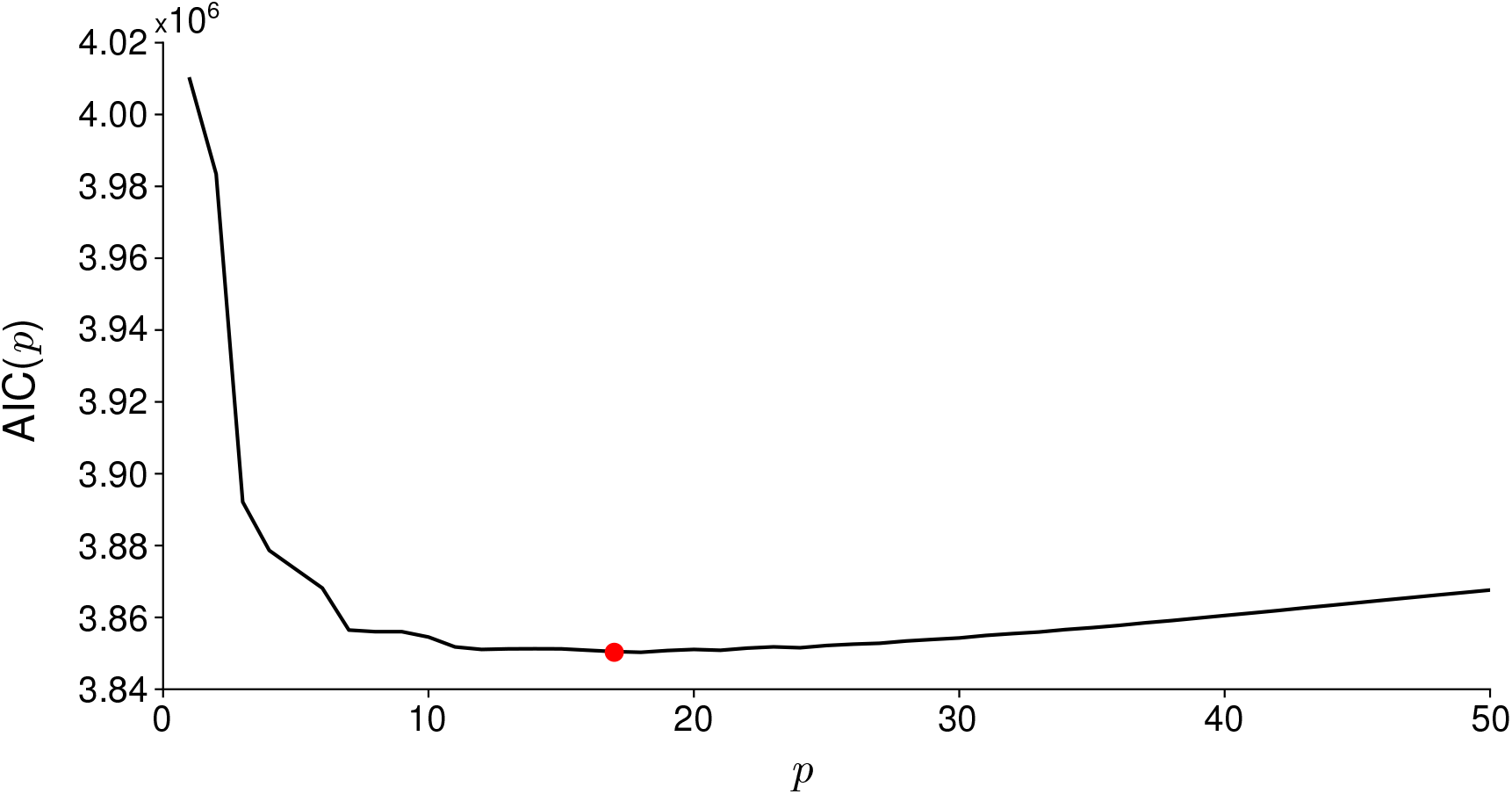
AIC for the analysis of one simulation. The best order (red bullet) is 17 with AIC(17) ≈ 3850286. For all simulations analyzed in Figure 2 the best order was 17 where the average AIC(17) over 10 simulations was 3855348.8 and the standard deviation was 4855.3.

**Table S2:** AIC for each cluster size. Minimum, median and maximum of distribution of AIC values for GPDC computed for each cluster size. The distribution consider GPDC computed for all simulations and all randomly chosen areas.

| [HTML]C0C0C0 Cluster size | Minimum AIC | Median AIC | Maximum AIC |
| --- | --- | --- | --- |
| 3 | 18 | 21 | 39 |
| 4 | 18 | 21 | 33 |
| 5 | 18 | 21 | 24 |
| 6 | 18 | 21 | 24 |
| 7 | 18 | 21 | 24 |
| 8 | 18 | 18 | 24 |
| 9 | 18 | 18 | 21 |
| 10 | 18 | 18 | 21 |
| 11 | 18 | 18 | 21 |
| 12 | 18 | 18 | 21 |
| 13 | 18 | 18 | 21 |
| 14 | 18 | 18 | 21 |
| 15 | 18 | 18 | 21 |

